# Who Are The Traditional Healers Treating Mental Illnesses In Rural Ethiopia? A population-Based Descriptive Study

**DOI:** 10.1101/543850

**Authors:** Ayele Belachew, Mitikie Molla, Abebaw Fekadu

**Affiliations:** Department of Preventive Medicine, School of Public Health, College of Health Sciences Addis Ababa University, Addis Ababa, Ethiopia; Department of Psychiatry, School of Medicine, College of Health Sciences Addis Ababa University, Addis Ababa, Ethiopia

**Author notes:** Corresponding author Ayele Belachew (AB) 251 913 306792.

**Keywords:** Ethiopia, Traditional and faith healers, Traditional practice, mental health care, mental illness.

## Abstract

**Background:** Ethiopia is a traditional country with a pluralistic health care system where people use the traditional health care systems in combination with the modern health care. In view of this, we assessed the profile of faith and traditional healers and their experience in treating people with mental illness in rural district Ethiopia so that inform the health minister to consider working towards integration with modern biomedical care to improve access.

**Methods:** We conducted a descriptive cross-sectional study among 173 traditional healers in Sodo district of Ethiopia. Data were collected through face-to-face interviews by trained enumerators using pretested structured questionnaires and analyzed using SPSS version 20.

**Result:** The median age of traditional and faith based healers was 55 (IQR=48.5, 67 years), about a third (29.5%) of them were female and 54(31.2%) earned their living exclusively from traditional healing practice. Eighty six (48.6%) healers didn’t attend formal education. Four types of healers were identified-herbalists accounted for 59% (n=102), faith healers were 36 (20.8%) mixed herbal and faith practitioners were 19(11.0%) and 16 (9.2%) were diviners. Most, 119(69%) had been practicing for an average of 15 years. Half of healers entered into the healing practice due to family kinship, whereas 26(15%) because of ancestral spirit.

Seventy one (41%) of the healers reported that they have ever treated patients with mental illness in their lifetime. Sixty three(36.4%) reported that they had treated mental illness within the past one year, of which 30(47%) treat only mental illness while 33(52%) treat both mental and physical illnesses. All faith healers and divine healers reported treating mental illness while 11(57.9%) of mixed healers, and no herbalists reported treating mental illness. Only 58(33.5%) believed that mental illness can be cured completely.

**Conclusion:** Significant proportion of traditional healers manages mental illness and remains an important part of the healthcare system in the rural setting of Ethiopia. Herbalists believed that biomedical treatments are preferable for mental illnesses, while faith healers and diviners believed traditional practices alone or in combination with biomedical practices is the treatment of choice.

## Background

Mental disorders are common health conditions and are associated with severe disability, cost and mortality. The burden of mental disorders in low income countries is compounded by the huge treatment gap, where in some countries, the lifetime treatment gap reaches 90% [1]. In these low income settings, 60%-80% of the population rely on traditional form of health care [2, 3]. A one year prevalence of use of traditional healers services was found to be 61% among South Africans [4]. In Ghana, 71% had consulted traditional healers in the previous one year while only 53% consulted modern care system [5]. There is a similarly high use of traditional services in Ethiopia [6].

Among the reasons that attract people to traditional care are cultural acceptability, relatively lower cost, accessibility [6, 7] availability, shared social norms, and beliefs about the meaning, cause and treatment of illness people share with Faith and Traditional Healers (FTHs) [7–9]. Additional factors considered relevant for choosing FTHs are simplicity and convenience [10], inclusiveness [11] and positive personal or family experience [12].

People become aware of FTH services and seek help from them by word of mouth, advice from friend and families, referral from fellow traditional healers as well as modern health care providers [13]. People are either self-referred or referred by someone else, including cross-referral between traditional healers [12]. People use traditional healing practices for all kinds of health problems including mental disorders, either exclusively or along with modern care concurrently or sequentially [13]. Peoples’ belief systems and culturally specific explanations of illness influence their acceptance of the care and support being provided [14]. People tend to attribute mental illness to supernatural phenomenon, and FTHs are viewed as having the expertise to address the illness than biomedical practitioner [15, 16].

Based on their primary practice FTHs can be divided in to four types: Faith healers (practicing prayer, recitation or sprinkling of holy water in churches, holy water places and mosques) use the power of God to heal sickness [17]; Divine healers (Ritualist or spiritualists, *kalechas, and Tenquay/wizard* in the case of Ethiopia, who practice astrology, read zodiac sign, etc) [18]; Herbalists (called *secular* healers and treat patients using herbs, plant remedies or even extracts of animal origin) [19]; and Mixed (Herbalist-Ritualist i.e. healers who use both rituals and herbal medicine). Faith healers and diviners are also called *spiritual* healers. In the Ethiopian context, *secular healers* are those involved in manipulation of body using a variety of techniques such as bone-setting by the *wegesha* (a physiotherapist/orthopedic surgeon) or assisting births by the *Yelimd awalaj* (traditional mid-wife), dressing wounds or excising affected body parts, draining abscesses, pulling out ‘bad’ teeth (tooth extractors), or cutting out the uvula or tonsils, inoculation, and provision of remedies such as herbs, minerals, animal products and thermal waters by the *medhanit awuqi* (herbalist) [20, 21]. *Spiritual healers* are the ones who claim that they have certain magical power, usually governed by either bad spirit to make a person ill or good spirit to protect form developing a psychotic illness, and included in this group are *at’maqi* (baptizer-exorcist), *Debtera* (cleric-diviner-healer) [20] ’*Tenquay*’ (witch doctors), ’*Bale-Weqaby*’ and ’*Kalicha*’ [22, 23]They exorcise malign spirits such as *buda* (evil eye) and *ganen* (devil) [20]by use of incantation, sorcery, enchantment and certain rituals [24]. Exorcism is a procedure that involves conducting special ceremonies, burning incense, praying, using holy water, and advise to put amulets containing a written script [16]

Almost all traditional practices are private practices and are entirely financed by patients or caregivers. Fees vary greatly and usually affordable by majority of the people, and can be made either monetarily or in kind. Traditional healers are an integral part of the society and the people, and are believed to share similar beliefs and attitude towards all life events (health and ill-health) as well as expectations with the people they are living [25].

In general, the number of traditional healers outweigh that of medical practitioners [2, 3, 10, 17, 26, 27], however, it is usually difficult to exactly know their number and type of their practice [28].

The agenda of establishing collaboration between the traditional and modern health services has long been advocated by the World Health Organization (WHO) to improve access to mental health care [29, 30]. Understanding FTHs in terms of basic socioeconomic and demographic characteristics, modalities of healing practices (what and how they are practicing), users’ physiognomies and healers explanatory models of mental illness as well as their attitude towards bio-medical service and intention to collaborate helps the development of evidence informed strategies to improve access to timely bio-medical care for Seever Mental Disorders (SMDs). Although traditional healers are an important part of Ethiopia societies, information about the extent and characteristic of healing and the people involved in the practice is limited [30]. This knowledge is important first step to introduce interventions as well as scale up of mental healthcare.

The main purpose of this study was to describe the profile of FTHs in a rural district in Ethiopia and their experience in managing mental illnesses. The study is part of the Programme for Improving Mental health care (PRIME), a study which works in this rural district to develop evidence on the best approaches of integrating mental health care [31, 32].

## Methods

A descriptive cross-sectional community based quantitative study was conducted among 173 faith and traditional healers found in Sodo district of Ethiopia, to describe who they are, their healing practices and treatment experiences in for mental illness. Sodo district is predominantly rural inhabited by Orthodox Christians and farmers. Study participants were all herbalists (those who use herbs, plant remedies or even extracts of animal origin to treat their patients), all diviners [spiritualists or who use rituals, *debteras and kalechas* (those who have some church education and defect from church services) *Tenquay* (wizard/ witch)], who practice astrology, read zodiac sign, etc), all faith healers (practicing specific healing prayer or recitation in church and in mosques, sprinkling of holy water at holy water places) and mixed, those who mix the above methods.

Participants were identified though household censes where names and the predominant type of traditional healing practices of all FTH (n=182) were listed using health extension workers three weeks prior to the actual study.

### Instrument

Data was collected through face-to-face interview using a pretested structured questionnaire, developed in English then translated in to their local language. Information about basic socio-demography, type of healing practices, clients’ characteristics, healing practices for mental illness, and experience and intention of FTHs to collaborate with modern health care system were collected.

### Data management and analysis

Statistical package for social sciences (SPSS) version 20.0 was used for data entry, cleaning and analysis. Descriptive statistics were done to summarize the profile of FTHs. Response frequencies were analyzed using Chi-square test at bivariate level to test for any association with treatment practices, experience of collaboration with modern care, and other important variables by type of healers whenever possible.

### Ethical consideration

Ethical clearance and approval of the study was obtained from the Addis Ababa University, College of Health Sciences, School of Public Health, Research Review Committee. All patients were fully informed about the purpose, benefit and harm if any associated with the study, and verbal informed consent was obtained. It was emphasized that voluntary participation was required and participants were informed their full right to refuse at any time to stop their participation and were assured their refusal to participate will not have any negative consequences.

## Results

### Socio-demographic characteristics of participants

We obtained a response rate of 95% where 173 FTHs participated in the study. Non-Muslim faith healers were interviewed at the healing sites (churches and holy water places) while Muslim faith healers, herbalists and diviners were interviewed in their residential places, where they usually provide healing services.

One hundred twenty two (70.5%) of the healers were male and the difference between the two sex was significant (χ^2^ test 13.05, df=3, p=0.01). The median age of healers was 55 years (IQR= 48.5, 67 years), and had lived in the locality for a median duration of 45 years (IQR= 35, 59 years). Greater majority 145(83.8%) were married and 158(91.3%) followers of Orthodox Christianity (91.3%). Fifty four (31.2%) earned their living exclusively from the FTH practice, while the rest had other work for additional income. Eighty four (48.6%) of healers didn’t attend formal education. (Table 1)

**Table 1:**
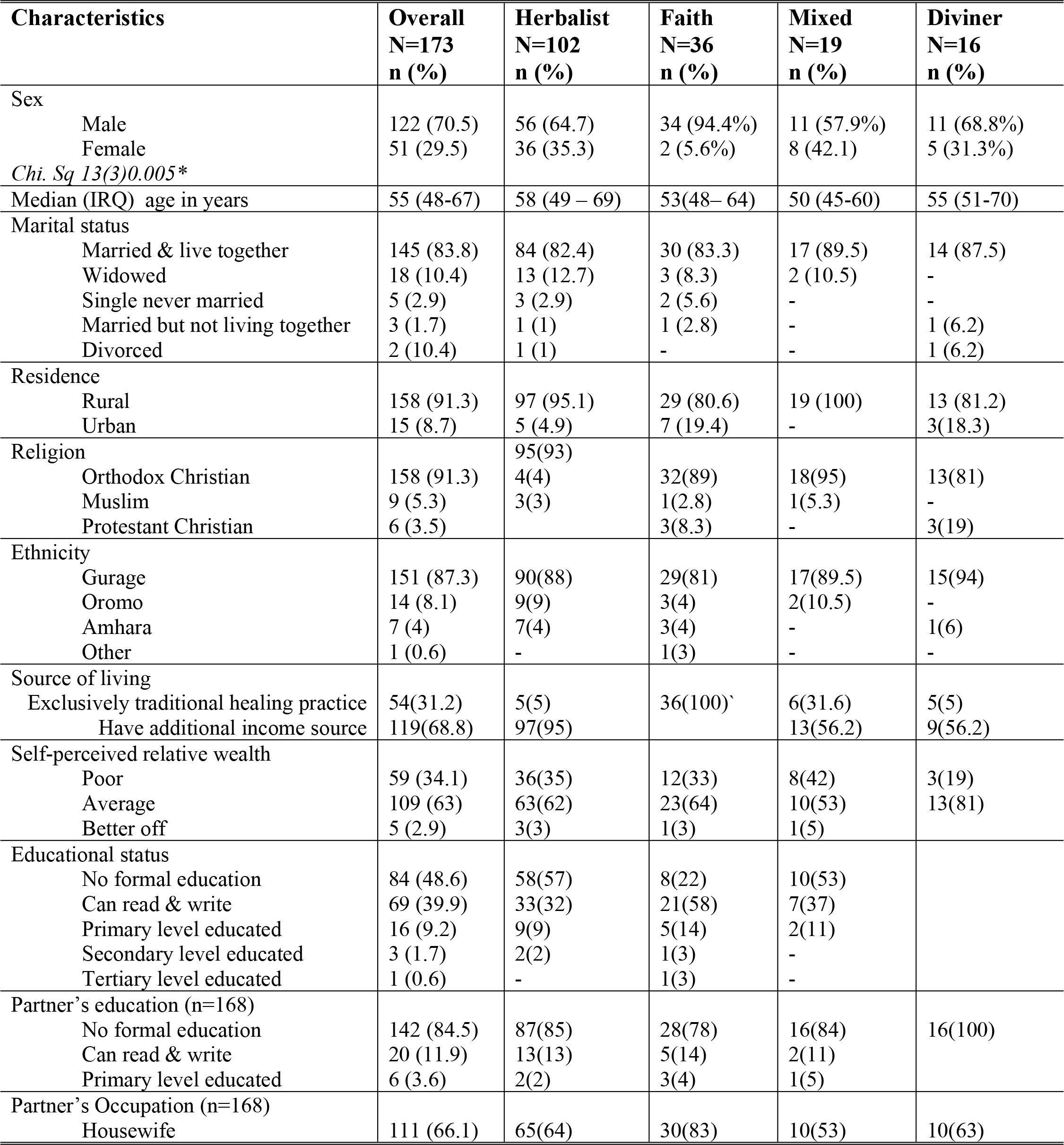
Socio-demographic and economic characteristics of FTHs, Sodo district, SNNPR, Ethiopia, 2014 (N=173)

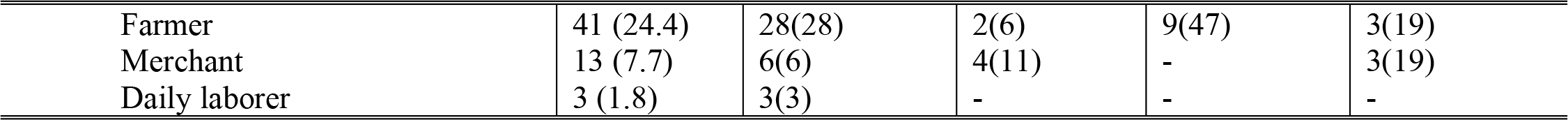

### Traditional healing practice

Four types of healers were identified: 102(59%) were herbalists, 36 (20.6) faith healers, 19 (11%) were mixed healers and 16 (9.2%) were diviners. About two-third (n=68) had history of illness, mostly physical illness, prior to becoming healers. The median years of practicing as healer was 25 years (IQR =13; 34 year). Ninety one (52.6%) of FTHs entered the healing practice through family kinship experiences and 36(20.9%) entered the practice because of their satisfaction with treatment they received from FTHs in the past. Only 15% entered the healing practice because of ancestral spirit. Very few FTHs (5.8%) received training before joining the healing practice. The training was generally very brief, on average two and half days. Training focused on how to cast out possessing spirits, on how to identify and apply herbs, and on how to select versus of the Bible which are amenable to treating mental illnesses. Ten (5.8%) of healers entered the practice after instructions by supernatural powers.

### Cost and utilization of services

Seventy four (42.7%) of healers receive payment for their healing service, of which 56(75.7%) accept payment in kind. The amount of payment varies, in such a way that 24(32.4%) said payment depends on the capacity (wealth) of the client/caregiver, while 20(27.3%) and 15(20.3%) said it is based on the type or severity of illness presentation, and on the desire of the client/caregiver respectively. On the other hand the amount was fixed as reported by 15(20.3%) of healers. The usual payment received for traditional healing service ranges from 2.3 USD to 6.65 USD per patient, and this amount was believed by healers as cheap compared to biomedical care.

In a typical busy day the average number of clients ranged from two to 20 patients. Majority of healers (96%) provide healing service every day whenever they were needed, and did not have specific preferred day or time of day dedicated for providing service. Overall, 54(31%) of FTHs have space for patients for overnight stay during treatment; however, all faith healers have accommodation for patients.

### Characteristics and source of clients

Traditional healers’ client were not distinguishable by gender, age, wealth or educational background, except residence. Majority (71.7%) of the healers receive clients from the same district that they are living in. One hundred nine (63%) of the healers were certain that people visit both traditional and modern treatments simultaneously and 100(57.8%) believed that patients preferred to visit biomedical care first. Almost all (95.4%) mentioned that their clients came to know them by what they have heard about them in the community (word of mouth) and 98(56.6%) also reported that family members advised patients to see healers.

When looking at who influenced patients to come to traditional healers, 102(59%) of healers believed that patients them without anyone’s influence, while 99(57.2%) and 81(46.8%) believed that patient’s family and friends or other person who received treatment by healers were main influencers to receive traditional treatment respectively. With regard to perceived reasons why people use traditional healing, 85(49%) believed that it is because of the availability of healers whenever they are needed, while 71(41%) and 62(35.8%) believed it is because healers existed for long time in the locality and the healers are more effective respectively (Table 2).

**Table 2:**
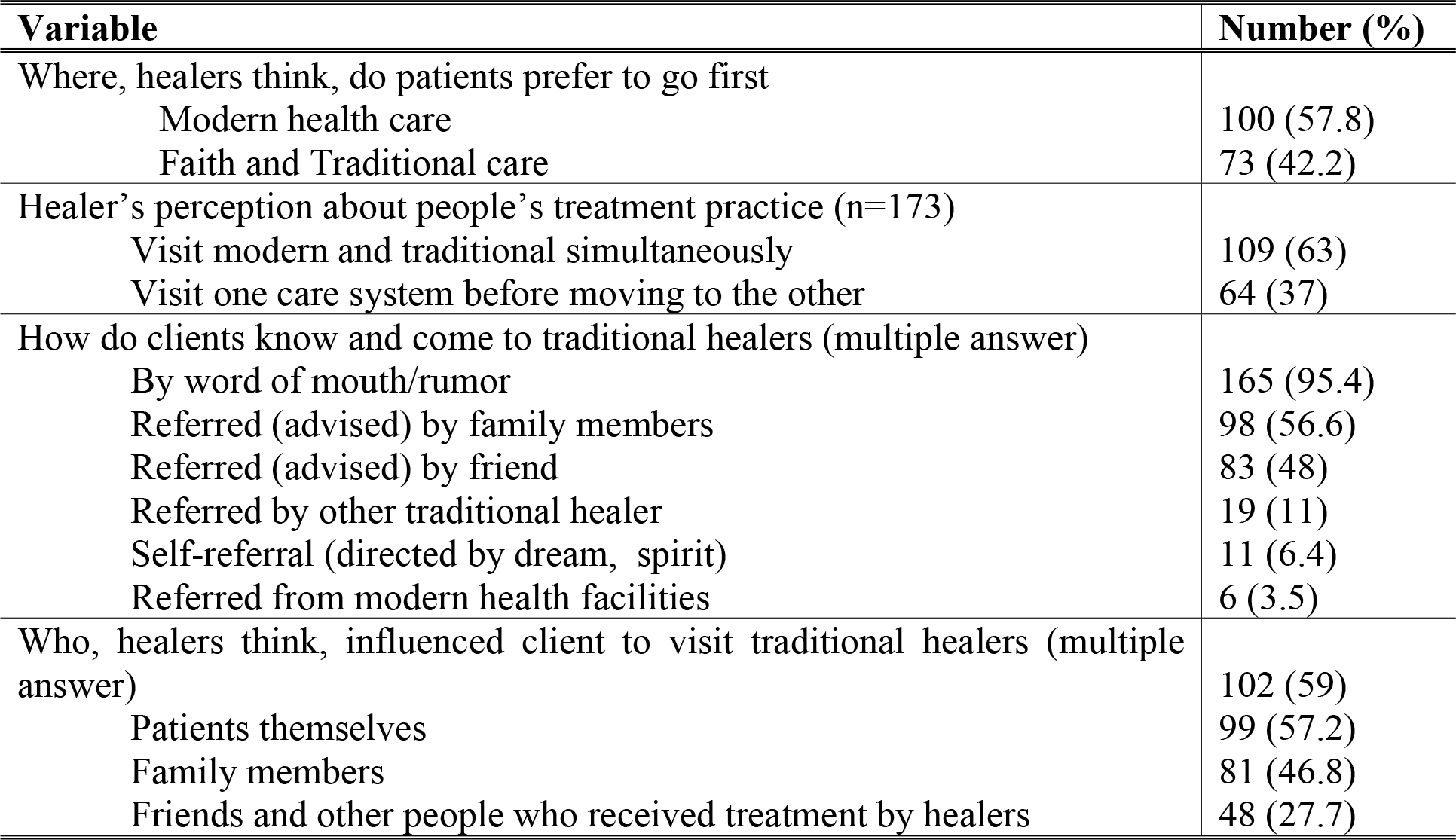
Source of clients and factors influencing use of service *Sodo district, SNNPR, Ethiopia, 2014 (N=173)*

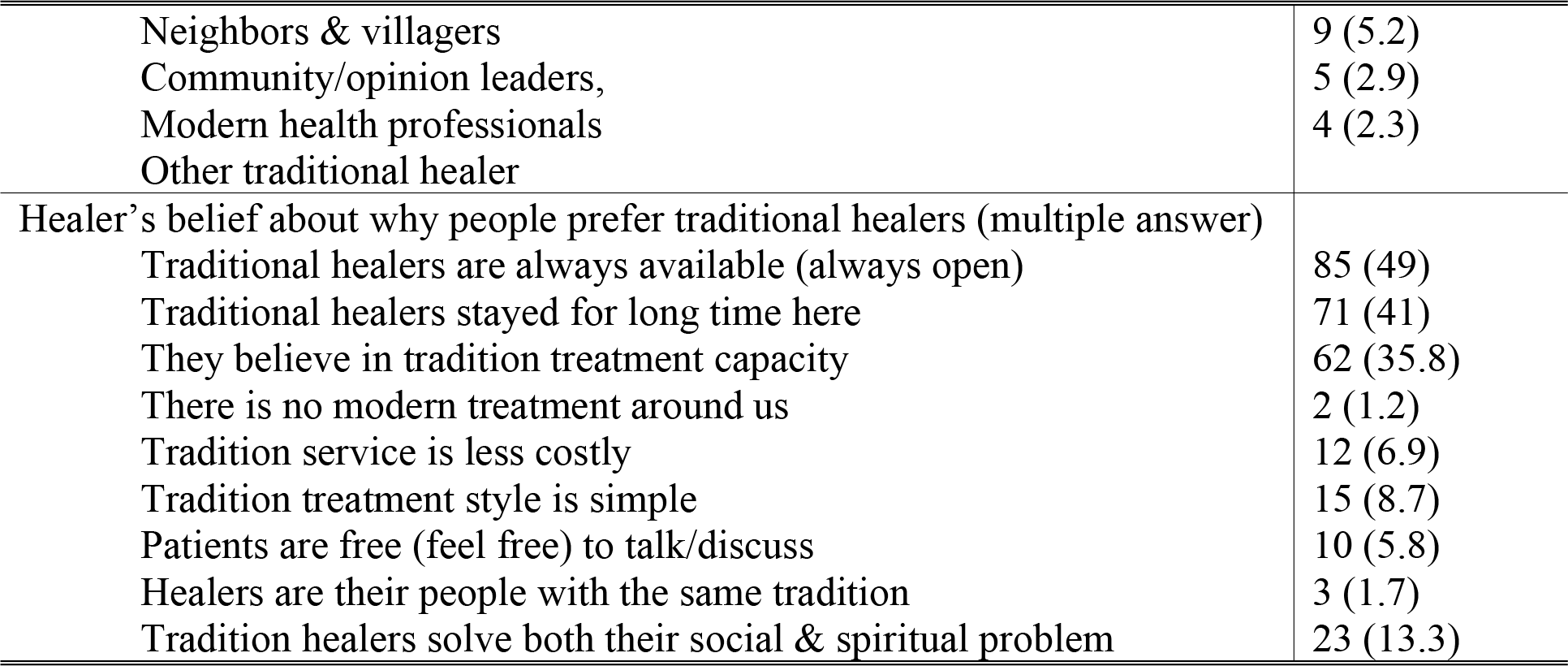

### Healer’s understanding about mental illness

Only 58(33.5%) of FTHs believed mental illness was common in their locality while 102 (59%) believed that it is not common, and the rest were no sure. With regard to altered behavioral manifestations in patients with mental illness, 96(55.5%) of healers cited talking alone or nonsense, 82(47.4%) mentioned shouting and 79(45.7%) and 71(41%) mentioned impaired self-care and laughing at inappropriate time or cause respectively. Concerning cause of mental illness 141(81.5%) mentioned supernatural causes. In a multiple response question, 109(63%) mentioned economic problem, while 43.1% and 42.2% cited curse and shock due to unfavorable life event (death, sudden loss of property, etc), respectively (Table 3).

**Table 3:**
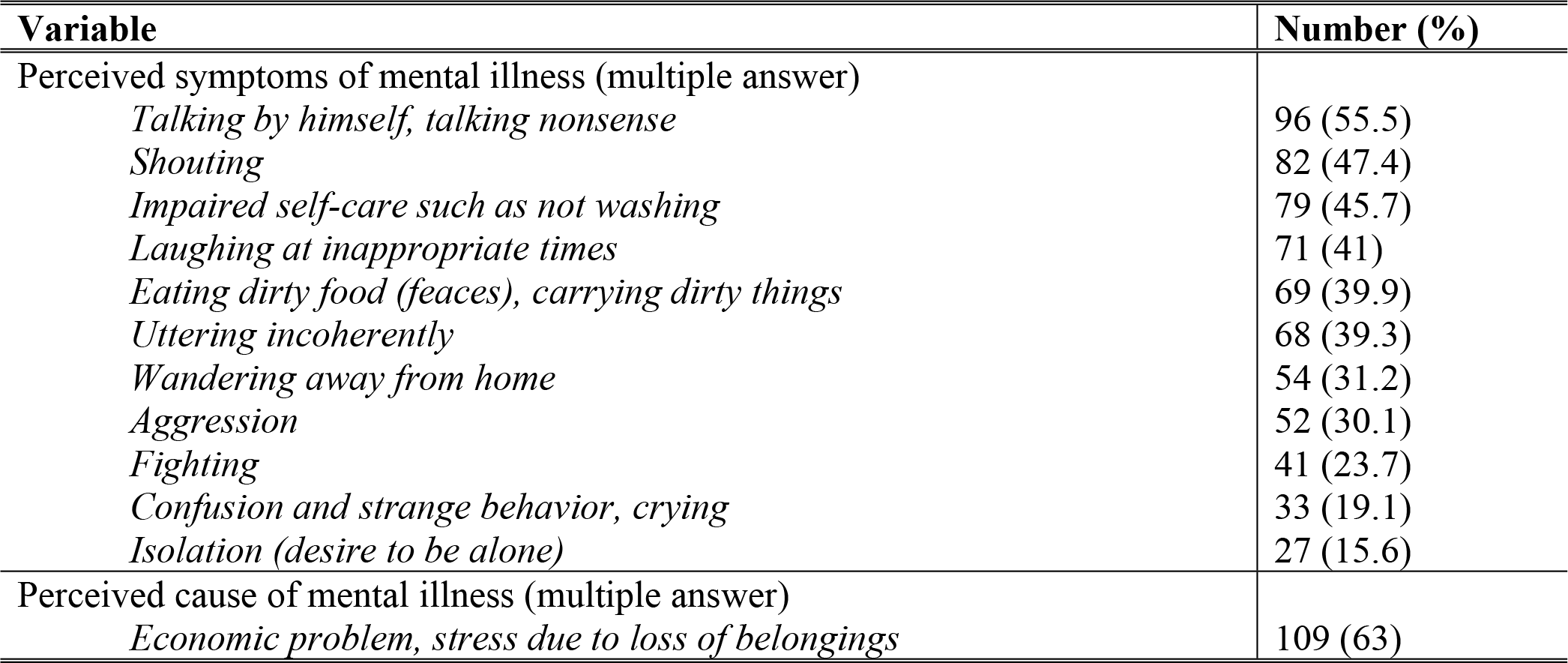
Perceived manifestations and causes of mental illness by FTHs, Sodo district, SNNPR, Ethiopia, 2014 (N=173)

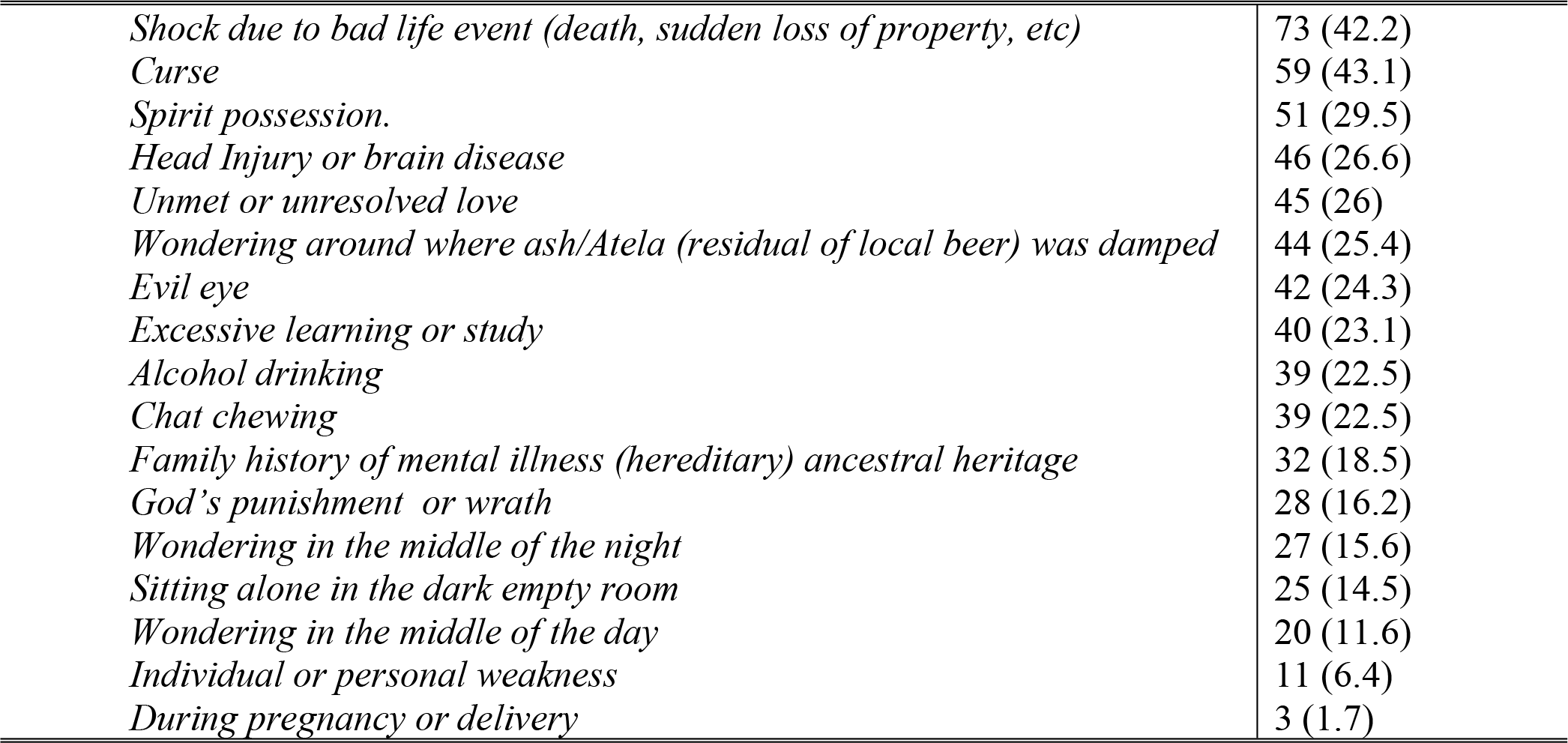

### Treatment practice for mental illness

Overall, 63(36.4%) reported experience of treating mental illness, of which 30(47%) treat only mental illness while 33(52%) treat both mental and physical illnesses. When looking by type of healers all faith healers and divine healers reported treating mental illness while 11(57.9%) of mixed healers, and no herbalists reported treating mental illness. On average in a typical month healers reported to treat two mentally ill patients.

With regards to prognosis of mental illness, only 58(33.5%) believed that it can be cured completely while 115(66.5%) believed no permanent cure only temporary improvement. Those healers who treat mental illness were asked about possible factors that influence the outcome of their treatment, and 40(32%) mentioned severity of illness (i.e., the strength or power of the spirit that possessed the patient), 42(24.3%) cited compliance to treatment, 36(20.8%) mentioned patient’s good general health and nutritional status while 34(19.7%) reported duration of illness.

## Discussion

In this study we found that, three of the four types of traditional healers (faith-based, diviners and those who mix herbal and faith) played an important role in the treatment of mental illnesses in the District. While those who do not treat mental illnesses (herbalists) believed that biomedical sciences is important in the treatment of mental illnesses. The belief that biomedical sciences could not be helpful for mental illnesses was also supported by the fact that almost all except herbalists indicated that the treatment outcome of mental illnesses depends on the power of the spirit that possessed the mentally ill person. Most Faith healers indicated their intention towards linking with biomedical care is a positive indication for improvement of the health care provided to mentally sick people. However, though few some had indicated that they have received patents referred by modern health care providers. This has boosted the morale of the FBTHs and confirmed their belief that modern health care is not able to treat mentally ill patents. This could be one of the threats that could affect linkage of the two health care systems. Traditional and faith based healers in the district indicated that they do not have any formal training but kinship and experience.

Traditional healing providers were widely available in this traditional and rural district. Herbalists were the most common traditional practitioners followed by faith healers. The vocation was dominated by men as appears to be the case in other African countries [17]. All the four types of FTHs share similar socio-demographic characteristics with each other and, for some domains, with the population of the district [31]. Most healers were also middle aged consistent with other studies conducted in Ethiopia [34]. These demographic characteristics are likely to have bearing on the service being provided. While the shared characteristics may indicate that traditional healers and the community may share explanatory models, other characteristics, for example age and gender differences may limit choice for patients; particularly, the service may not be equitable, acceptable or appropriate for women. Further exploration of the impact of FTHs characteristics and how they may be perceived by women and other groups in the locality may be important in pursuing collaborative care and improving accessibility of care.

Family kinship is an important path of entry in to the healing practice; however, experience of illness followed by healing by healers was also another important path to the practice. These pathways were reported by a similar study from Uganda [28]. More interesting was the fact that very few healers had received any kind of training or mentorship from anywhere when they started the healing practice. The impact of this lack of training is unclear and is worth exploring further.

A relatively large number of service users reportedly use the traditional services every day. On average, about 415 clients visit traditional healers daily. This is much larger than the number that would be receiving care daily across the eight biomedical centres in the district, which is about 160. This makes the traditional healing services the “de facto” primary care for the people in the Sodo district. This is also unlikely to be much different for other parts of the country. Geographic, cultural and economic accessibility is likely to be at the heart of this healers’ service utilization.

The situation is not likely to change in the immediate feature and collaborating with traditional healers is essential. Yet, virtually no collaboration exists between traditional and biomedical services. Although some traditional healers believe they have the better ability to cure illnesses, some blame biomedical providers for ignoring the service of traditional healers. Whatever the role of traditional healers might be, services to the people will be accessible only when collaboration is strengthened and when biomedical services are available more widely. The fact that most users come to traditional healers through word of mouth and advice from family or friend does also suggest that traditional healers are recognized, valued and trustworthy providers. Biomedical providers have to be similarly trustworthy by residents if they are to be useful and more accessible. It is also of note that availability of the service at all times and the ability of healers to solve social and spiritual problems of the people [7–9] were considered important assets by the traditional healers of Sodo. These are important lessons for biomedical providers. Providing holistic care, care that is available all the time in close proximity to the people is more likely to be used and possibly trusted by the people.

According to the report of FTHs, users of the FTH servicers do not differ by their wealth, unlike in South Africa where majority of users belong to the poorer sections of the society [17], and perhaps our study population were different from the South Africans with respect to the general economic standard, and being rural as well. Furthermore, reported users do not differ by sex and educational status unlike users in the United States of America where women than men, educated than less educated were more likely to use healing [35]. However, actual data from users is required to make judgment about the characteristics of users. It was against our assumption that only a small minority of healers provided care for mental illness.

## Conclusion

The study is the first comprehensive study looking at the role of faith and traditional healers in the care of mental illness in Ethiopia. Traditional practices are the “de facto” primary care providers in this rural district. All faith healers and diviners providing care for mental disorders though few herbal medical practitioners do so. It is essential to involve traditional healers in any engagement to improve access to care. We recommend a wider community based mixed qualitative and quantitative study both on traditional healers and clients to document the situation in depth.

## Data availability statement

Data are available within the Supporting Information files, and can be submitted upon request

## Acknowledgments

Authors acknowledge the Addis Ababa University and those faith and traditional healers who consented and participated in the study.

## List of abbreviations

df: Degree of Freedom
FTHs: Faith and Traditional Healers
FTHp: Faith and Traditional Healing Practice IQR Inter Quartile Range
SMDs: Sever Mental Disorders
χ^2^: test Chi Square
SNNPR: Southern National Nationalities and Peoples Region WHO World Health Organization

## Competing interests

The authors have no competing interests.

## Authors Contribution

**Conceptualization:** Ayele Belachew Aschalew,

**Data collection**: Ayele Belachew Aschalew

**Formal analysis:** Ayele Belachew Aschalew, Abebaw Fikadu

**Funding acquisition**: Ayele Belachew Aschalew, Abebaw Fikadu

**Investigation and measurements:** Ayele Belachew Aschalew, Abebaw Fikadu

**Methodology:** Ayele Belachew Aschalew, Abebaw Fikadu

**Supervision:** Ayele Belachew Aschalew, Abebaw Fikadu

**Writing draft**: Ayele Belachew Aschalew, Abebaw Fikadu, Miteke Molla

**Review & Editing:** Ayele Belachew Aschalew, Abebaw Fikadu, Miteke Molla

